# Control of Transposon-mediated Activation of the *glpFK* Operon of *Escherichia coli* by two DNA binding Proteins

**DOI:** 10.1101/046649

**Authors:** Zhongge Zhang, Milton H. Saier

**Affiliations:** Department of Molecular Biology, Division of Biological Sciences, University of California at San Diego, La Jolla, CA 92093-0116.

**Keywords:** stress-induced mutagenesis, transposon, glycerol utilization, nutrient starvation, cyclic AMP-Crp

## Abstract

*Escherichia coli* cells deleted for the cyclic AMP (cAMP) receptor protein (Crp) gene (Δ*crp*) cannot utilize glycerol because cAMP-Crp is a required positive activator of glycerol utilization operon *glpFK*. We have previously shown that a transposon, Insertion Sequence 5 (IS*5*), can reversibly insert into the upstream regulatory region of the operon so as to activate *glpFK* and enable glycerol utilization. GlpR, which represses *glpFK* transcription, binds to the *glpFK* upstream region near the site of IS*5* insertion, and prevents insertion. We here show that the cAMP-Crp complex, which also binds to the *glpFK* upstream regulatory region, also inhibits IS*5* hopping into the activating site. This finding allowed us to identify conditions under which wild type cells can acquire *glpFK*-activating IS*5* insertions. Maximal rates of IS*5* insertion into the activating site require the presence of glycerol as well as a non-metabolizable sugar analogue that lowers cytoplasmic cAMP concentrations. Under these conditions, IS*5* insertional mutants accumulate and outcompete the wild type cells. Because of the widespread distribution of glucose analogues in nature, this mechanism of gene activation could have evolved by natural selection.

## Introduction

Wild type *E. coli* cells can grow on glycerol as a sole carbon source, but cells lacking the cAMP receptor protein (Crp) cannot (Lin, 1976, Won et al., 2009, Fic et al., 2009). In a previous communication (Zhang and Saier, 2009a), we showed that a Δ*crp* strain could mutate to rapid glycerol utilization due to insertion of the small transposon, Insertion Sequence *5* (IS*5*) (Sousa et al., 2013). To cause activation, IS*5* hops into a single site, in a single orientation, upstream of the *glpFK* operon promoter. The presence of IS*5* at this site activates the *glpFK* promoter so that it becomes stronger than that in wild type cells (Zhang and Saier, 2009a). Interestingly, a gene-activating IS*5* insertion is capable of undergoing subsequent precise excision (Zhang et al., 2010), showing that IS*5*-mediated operon activation can be a fully reversible process. The *glpFK*-activating insertional event occurred at high frequency in the presence of glycerol, but not in the presence of glucose or other carbon sources. Glycerol increased insertion of IS*5* at this specific site, but not in other operons (Zhang and Saier, 2009a, Zhang and Saier, 2011). Glycerol-promoted IS*5* insertion into the *glpFK*-activating site proved to be regulated by binding of the glycerol repressor, GlpR, when bound to the four adjacent *glpFK* operators, *O1, O2, O3* and *O4* in the *glpFK* control region. However, it became clear that the effect of GlpR-binding on IS*5* insertion was not mediated by increased expression of *glpFK*, or by increased growth, since binding to *O1* primarily controlled IS*5* insertion without a significant impact on transcription, while binding to *O4* primarily controlled transcription (Zhang and Saier, 2009a). Thus, the negative control of IS*5* insertion into the upstream activating site is a newly recognized function of GlpR that is distinct from the previously recognized function of repressing *glpFK* transcription (Zhang and Saier, 2011).

In a previous study, Hall has summarized situations where transposon insertion to activate cryptic operons is elevated under conditions of starvation (Hall, 1999). More recently, Wang and Wood (Wang and Wood, 2011) described another example of IS*5* insertion that occurs at a higher frequency when beneficial, giving rise to activation of the *E. coli* flagellar master switch operon, *flhDC*. Insertional activation of *flhDC* substantially enhances bacterial swarming in semisolid agar media, and the swarming phenotype is beneficial under adverse conditions such as nutrient depletion. Under conditions where swarming is not permitted (liquid or on solid agar media), insertional activation of the *flhDC* operon is not beneficial, and the frequency of insertion is greatly reduced. Thus, insertion of IS*5* upstream of the *flhDC* operon is another example of a mutation whose frequency is elevated under conditions where the mutation is advantageous. We have now confirmed the observations of Wang and Wood, 2011, in our laboratory (Zhang et al., 2013).

Since *de novo* IS*5* insertion at a single site in the *glpFK* operon in Δ*crp* cells is promoted by starvation in the presence of glycerol, and the insertion event relieves the starvation by enabling glycerol utilization, the question arises whether this constitutes an evolved mechanism. However, in wild type (*crp^+^*) cells, under the tested laboratory conditions, no such IS*5* insertion mutants were obtained (Zhang and Saier, 2009a, Herring et al., 2006, Cheng et al., 2014). Our earlier studies had been conducted in a Δ*crp* deletion strain which grows poorly and consequently is not present under normal environmental conditions.

In this communication, we first report that in Δ*cyaA crp*^+^ cells, IS*5*-mediated *glpFK* activation occurs in a manner strictly analogous to that observed in Δ*crp* mutant cells. The *cyaA* gene codes for the cyclic AMP biosynthetic enzyme, adenylate cyclase, Cya (Gancedo, 2013). We further show that addition of cAMP to the growth medium, known to increase the cytoplasmic cAMP concentration (Saier et al., 1982), greatly suppressed IS*5* insertion specifically to this site. This effect occurred independently of GlpR, but it depended exclusively on Crp and the two adjacent Crp binding sites (CRPI and CRPII) that overlap the two GlpR binding sites *O2* and *O3*, in the *glpFK* control region (Zhang and Saier, 2009a, Weissenborn et al., 1992). It thus became clear that the conditions that predispose the *glpFK* operon to activation by IS*5* in wild type cells were (i) the presence of glycerol, and (ii) the presence of an environmental agent that could lower cytoplasmic cAMP levels.

Non-metabolizable glucose analogues and other sugar substrates of the phosphoenolpyruvate (PEP):sugar phosphotransferase system (PTS) are among the compounds known to lower cellular cAMP concentrations by inhibiting adenylate cyclase (Feucht and Saier, 1980). These include 2-deoxy-D-glucose (2DG) and methyl-α-D-glucoside (αMG) (Saier et al., 1982). Here we show that incubation of wild type *E. coli* cells in glycerol media together with 2DG or αMG promotes *glpFK-*activating IS*5* insertional events. To determine whether under these conditions IS*5* mutants can accumulate and take over the population, we carried out long-term growth experiments. Our results are consistent with a scenario in which IS*5* insertional activation of *glpFK* evolved in wild type cells as a mechanism to overcome adverse environmental conditions.

## Materials and Methods

### Bacterial Strains and Growth Conditions

Strains and DNA oligonucleotides used in this study are described in Supplementary Tables 1 and 2, respectively. The *cyaA* deletion mutant was generated from the parental strain (*E. coli* K-12 strain BW25113) using the method of Datsenko and Wanner (Datsenko and Wanner, 2000b). Briefly, a kanamycin resistance gene (*km*), flanked by the FLP recognition site (FRT) was amplified from the template plasmid pKD4 using mutation oligos cyaA1-P1 and cyaA2-P2 (Supplementary Table 2), each of which is composed of a ~20 bp region at the 3’ end that is complementary to the FRT-flanking *km* sequence, and a ~50 bp region at the 5’ end that is homologous to *cyaA*. The PCR products were gel purified, treated with *Dpn*I, and then electroporated into BW25113 cells expressing the lamada-Red proteins encoded by plasmid pKD46. The pKD46 plasmid, which carries a temperature-sensitive origin of replication, was removed by growing the mutant cells overnight at 40 ^o^C. The Km^r^ mutants were verified for the replacement of the target gene by the FRT-flanking *km* gene by PCR. The *km* gene was subsequently eliminated (leaving an 85-bp FRT sequence) using plasmid pCP20 that bears the FLP recombinase. The *cyaA glpR* double mutant was constructed by transferring a *km* insertional mutation of the *cyaA* gene into the *glpR* deletion mutant background (Zhang and Saier, 2009a) using P1 transduction.

To fuse the chloramphenicol-resistance gene (*cat*) with the *glpFK* operon, downstream of *glpK* in the chromosome, the plasmid pKD13-*cat* made previously (Zhang and Saier, 2009a), was used. In this plasmid, the *cat* gene is located upstream of a FRT-flanking *km* gene (Datsenko and Wanner, 2000b). The *cat* structural gene with its own ribosome binding site (RBS), together with its downstream *km* gene, was amplified from pKD13-*cat* using primers *glpFK*cat1-P1 and *glpFK*cat2-P2 (Supplementary Table 2). The PCR products were electroporated into wild type, Δ*cyaA* and Δ*cyaA* Δ*glpR* cells to replace the 85-bp downstream region between the 8^th^ nucleotide and the 94^th^ nucleotide relative to the *glpK* stop codon in the chromosome. After electroporation, the cells were selected on LB + Km agar plates. The Km^r^ colonies were verified for the substitution of the 85 bp *glpK*/*glpA* intergenic region by PCR and subsequent DNA sequencing. In the resultant strains (named BW_Cat, Δ*cyaA*_Cat and Δ*cyaA* Δ*glpR*_Cat, respectively), *glpF, glpK* and *cat* form a single operon with its expression solely under the control of the *glpFK* promoter (P*glpFK)*.

Strains were cultured in LB, NB or minimal M9 media with various carbon sources at 37ºC or 30ºC. When appropriate, kanamycin (Km; 25 μg/ml), ampicillin (Ap; 100 μg/ml), or chloramphenicol (Cm; 16-60 μg/ml) was added to the media.

### Mutations of Chromosomal Crp Operators

To modify the chromosomal Crp binding sites in the control region of the *glpFK* operon, the previously made plasmid pKD13-P*glpFK* (Zhang and Saier, 2009a), was used. In this plasmid, P*glpFK* and the FRT-flanking *km* gene were oriented in opposite directions. Using the quick-change site-directed mutagenesis kit (Agilent) and oligos P*glpFK*_CrpI&II_-F and P*glpFK*_CrpI&II_-R (Supplementary Table 2), both Crp operators (O*_CrpI_* and O*_CrpII_*) in the *glpFK* control region, contained within pKD13-P*glpFK*, were mutated by changing tatgacgaggcacacacattttaagt (-69 to –44 relative to +1 of P*glpFK*) to <u>g</u>a<u>cag</u>cgaggca<u>tctg</u>cattttaa<u>tc</u> (substitutions are underlined). The substitutions were confirmed by sequencing. Using the resultant plasmid, pKD13-P*glpFK_O_CrpI&I I_*, as template, the region containing the *km* gene and P*glpFK_O_CrpI&II_* was PCR amplified using the primers P*glpFK*_CrpI&II_-P1 and P*glpFK*_CrpI&II_-P2 (Supplementary Table 2). The PCR products were integrated into the Δ*cyaA* mutant chromosome to replace the wild type P*glpFK*. The nucleotide substitutions in both O*_CrpI_* and O*_CrpII_* operators were confirmed by sequencing. The *km* gene was removed, and the resultant strains were named Δ*cyaA O_CrpI&II_* (Supplementary Table 1).

### Glp^+^ Mutation Assay Using a ΔcyaA Mutant Strain

Using the Δ*cyaA* deletion mutant, mutation to Glp^+^ was first measured on minimal M9 + 0.2% glycerol agar plates as described previously (Zhang and Saier, 2009a). Briefly, cells from an overnight LB culture were washed and inoculated onto plates (~10^8^ cells/plate). The plates were then incubated in a 30 ºC incubator and were examined daily for the appearance of Glp^+^ colonies with each colony representing a Glp^+^ mutation. On these glycerol minimal agar plates, any colonies appearing by day 2 were considered to be from Glp^+^ cells initially present when applied to the plates. They were therefore subtracted from the subsequent measurements. The total numbers of Glp^-^ cells were determined as described by Cairns and Foster (Cairns and Foster, 1991). The Glp^+^ mutations were determined by counting the Glp^+^ colonies that appeared on the original agar plates. The frequencies of Glp^+^ mutations on glycerol M9 plates were determined by dividing the numbers of Glp^+^ colonies by the total Glp^-^ populations. To determine if any of the Glp^+^ colonies arose from Glp^+^ cells initially plated, the Δ*cyaA* cells, together with small numbers of Δ*cyaA* Glp^+^ cells, were plated onto the same M9 + 0.2% glycerol plates. The plates were incubated and examined as above.

To determine the effect of cAMP on the frequency of IS*5* insertion into the *glpFK* activating site, strain Δ*cyaA*_Cat (in which *glpF, glpK* and *cat* are fused in a single operon, see Supplementary Table 1) was used. This strain is sensitive to Cm at 8 μg/ml while the same strain with the IS*5* insertion (Δ*cyaA* Glp^+^_Cat) is resistant to Cm at 16 μg/ml. Preliminary experiments showed that all Δ*cyaA*_Cat cells resistant to Cm at 16 μg/ml were due to IS*5* insertion in front of P*glpFK*. To determine the effect of cAMP on IS*5* insertion, an 8-h old culture from a single Δ*cyaA*_Cat colony was diluted 1000 x into 5 ml LB ± cAMP (0 to 5 mM) contained in 30 ml glass tubes (2.5 cm x 20 cm). The tubes were shaken at 250 rpm in a 30 ^o^C water bath shaker. After 15 h, the cells were washed 1x (to remove residual cAMP) with carbon source-free M9 salts, serially diluted, and applied onto LB + glucose agar plates and LB + glucose + Cm agar plates. The plates were incubated at 37 ^o^C for 15 to 18 h. Total populations and Glp^+^ populations were determined based on numbers of colonies on LB + glucose plates and on LB + glucose + Cm plates, respectively. The frequencies of Glp^+^ mutation were determined by the ratios of Cm^r^ populations to total populations.

To determine if cAMP affects IS*5* insertion into other chromosomal sites, we chose to analyze mutants resistant to Furazolidone (FZD) using Δ*cyaA*_Cat cells. First, step I mutants were isolated by spreading the cells onto nutrient broth (NB) agar plates with a low concentration (1 μg/ml) of FZD. The cells of a step I mutant were then applied onto the agar plates with higher concentrations (5-7.5 μg/ml) of FZD. The plates were incubated at 30 ^o^C for 36 h or above, and colonies obtained were examined for the presence of IS elements in the *nfsB* gene by PCR using oligos nfsB-ver-F and nfsB-ver-R (Supplementary Table 2). Among those mutants carrying IS elements, IS*5* insertional mutants were determined by two rounds of PCR, using oligos IS*5*-ver-F / nfsB-ver-R and nfsB-ver-F / IS*5*-ver-F, respectively. The ratio of IS*5* mutants was calculated by dividing total IS mutant numbers by IS*5* mutant numbers.

To establish the effects of mutations in the Crp operators in the *glpFK* control region on the appearance of Glp^+^ IS*5* insertional mutations, the Δ*cyaA O_CrpI&I I_*_Cat cells with mutations in operators, O*_CrpI_* and O*_CrpII_*, were examined for the appearance of Glp^+^ mutations in LB media with or without cAMP as described above. To determine the effect of loss of *glpR* on Glp^+^ mutation frequency, the Δ*cyaA* Δ*glpR* double mutant was examined for Glp^+^ mutations in liquid LB ± cAMP (0.1 mM) as compared to the single Δ*cyaA* mutant as described above.

To determine the effect of *glpR* overexpression on the Glp^+^ mutations, the *glpR* structural gene was amplified from the wild type genomic DNA using primers glpR-KpnI and glpR-BamHI (Supplementary Table 2). The PCR products were digested with *Kpn*I and *BamH*I, gel purified, and then ligated to the same sites of pZA31 (Lutz and Bujard, 1997), yielding pZA31-*glpR*, in which *glpR* is driven by a synthetic *tet* promoter (P*tet*). The same plasmid carrying a random fragment (RF), pZA31-RF, served as a control (Levine et al., 2007a). To repress P*tet* activity, the constitutively expressed *tetR* cassette, located at the *attB* site, was transferred to Δ*cyaA*_Cat carrying pZA31-*glpR* from BW-RI (Levine et al., 2007b) by P1 transduction. Therefore, the expression of *glpR* could be induced by a tetracycline analog, chlorotetracycline (cTc). The resultant strain containing the *tetR* source and pZA31-*glpR* was tested for Glp^+^ mutations in LB ± 0.1mM cAMP as described above. To induce expression of *glpR* in pZA31-*glpR*, cTc (250 ng/ml) was added to the medium.

To further demonstrate cAMP inhibitory effects on the appearance of IS*5* insertional mutants, and the competitive abilities of the mutants under low cAMP conditions, we performed a long-term experiment by transferring cultures to new media at various intervals. To do this, an LB culture (10 μl) of Δ*cyaA*_Cat was used to inoculate M9 + glycerol ± cAMP (0 to 1 mM) media. Before the first transfer, every 12 h or so, the cultures were serially diluted onto LB + glucose plates for total population determination and onto LB + glucose + Cm plates for Glp^+^ (IS*5* insertion) mutant population determination. After 2 or 2.5 days, the cultures were 1000x diluted into new tubes with the same media. Then at one-day intervals, the cultures were 1000x diluted into fresh media. For each transfer, the total cells and the Glp^+^ mutant populations were determined.

### IS*5* Insertional Mutation Assay Using A Wild Type Background

2-deoxyglucose (2DG) is a non-metabolizable glucose analog that reduces the level of cytoplasmic cAMP level when adding to the media (Saier, 1989). Preliminary experiments showed that *E. coli* cells are sensitive to this compound at 0.1%. To determine if IS*5* insertion upstream of P*glpFK* occurred in a wild type background, BW_Cat cells (Supplementary Table 1) were tested for IS^5^ insertion on M9 + glycerol (0.2%) + 2DG (0.13%) ± Cm (60 μg/ml) agar plates. The plates were incubated at 30 ^o^C, and the colonies were examined for the presence of IS^5^ in the upstream *glpFK* operon control region by PCR followed by gel electrophoresis.

### Long Term Evolutionary Experiments

At least two types of mutations arose when BW_Cat cells were incubated on M9 + glycerol + 2DG ± Cm agar plates, the IS^5^ insertional mutation and a non-IS^5^ mutation of unknown nature. To determine if the IS^5^ insertional mutants are more competitive than the non-IS^5^ insertional mutants, 10 μl of a fresh LB culture of BW_CAT was used to inoculate 5 ml of M9 + glycerol (0.2%) + 2DG (0.13%) ± Cm (60 μg/ml) in 30 ml glass tubes. The tubes were incubated with shaking (250 rpm) in a 30 ^o^C water bath shaker. After 2.5 days of incubation, the cultures (i.e., mutant cells resistant to 2DG and Cm) were 1000x diluted into new tubes with the same media. On every other day, the mutants were 1000x diluted into new tubes. For each transfer, the mutant cultures were serially diluted using carbon source-free M9 salts, and the 10^5^fold and 10^6^-fold dilutions were applied onto M9 + glycerol + 2DG ± Cm agar plates before incubation at 37 ^o^C. After 2 days, 100 colonies from each transfer were subjected to PCR and subsequent gel electrophoresis analyses to determine the percentages of IS^5^ insertion mutants to the total mutants. Note that the parental cells do not grow under these conditions.

### Chromosomal lacZ Fusions and β-Galactosidase Assays

Using pKD13-P*glpFK* (Zhang and Saier, 2009a) and pKD13-P*glpFK*_*O_CrpI&II_* (Supplementary Table 1) as templates, P*glpFK* (-204 to +66 relative to the transcriptional start site) and P*glpFK*_*O_CrpI&I I_* plus their upstream FRT-flanked *km^r^* gene were amplified using oligos P*glpFK*z-P1 and P*glpFK*z-P2 (Supplementary Table 2). Using the method of Datsenko and Wanner (Datsenko and Wanner, 2000a), the promoters plus the upstream *km^r^* gene (*km*^r^:P*glpFK* or *km*^r^:P*glpFK*_*O_CrpI&II_*) were integrated into the chromosome to replace the *lacI* gene and the native *lac* promoter (including the 5’ UTR of *lacZ*) of MG1655 deleted for *lacY* (Klumpp et al., 2009). This chromosomal replacement was confirmed by PCR and subsequent DNA sequencing analysis. The resultant strains are deleted for both *lacI* and *lacY*, but they carry the *lacZ* gene that is expressed under the control of P*glpFK* or P*glpFK*_*O_CrpI&II_*. Both constructs were transferred into Δ*cyaA* and Δ*cyaA* Δ*glpR* strains by P1 transduction, yielding Δ*cyaA*_P*glpFK*-*lacZ*, Δ*cyaA*_P*glpFK*_*O_Crp_*-*lacZ*, Δ*cyaA*Δ*glpR* _P*glpFK*-*lacZ*, and Δ*cyaA*Δ*glpR* _P*glpFK*_*O_Crp__lacZ* (Supplementary Table 1).

For β-galactosidase assays, strains were cultured in liquid LB media ± 1 mM cAMP at 30 ^o^C. When cultures entered the exponential phase, samples were collected for measurement of β-galactosidase activities as described by Miller (Miller, 1972).

## Results

### IS^5^ insertional activation of the glpFK operon occurs in ΔcyaA mutant cells

We previously demonstrated that Δ*crp* mutant cells of *E. coli* could regain the ability to utilize glycerol by IS^5^-mediated insertional mutations that specifically occurred at a single site, upstream of the *glpFK* promoter, preferentially in the presence of glycerol (Zhang and Saier, 2009a). In this study, we first used Δ*cyaA* mutant cells lacking the cAMP biosynthetic enzyme, adenylate cyclase, Cya. Like the Δ*crp* mutant cells, the Δ*cyaA* mutant cells cannot utilize glycerol as the sole carbon source for growth. However, after a prolonged incubation on M9 + glycerol agar plates, Glp^+^ mutants could be observed (Figure 1). These mutants were IS^5^ insertional mutants carrying IS^5^ in the same position and orientation upstream of the *glpFK* promoter as those isolated previously from Δ*crp* cells. The time course (Figure 1) for their appearance and the properties of these double mutants were indistinguishable from those isolated previously (Zhang and Saier, 2009a). Among at least 100 independent Glp^+^ mutants analyzed, no other types of mutants arose under the conditions used. There was an approximately two-day delay before mutant colonies appeared, and these clearly arose during incubation on the plates, since when small numbers (e.g. 11 and 23) of identical Δ*cya* Glp^+^ insertional mutants were added to the Δ*cyaA* cells prior to plating, they gave rise to colonies within a shorter time period (Figure 1).

**Figure 1.**
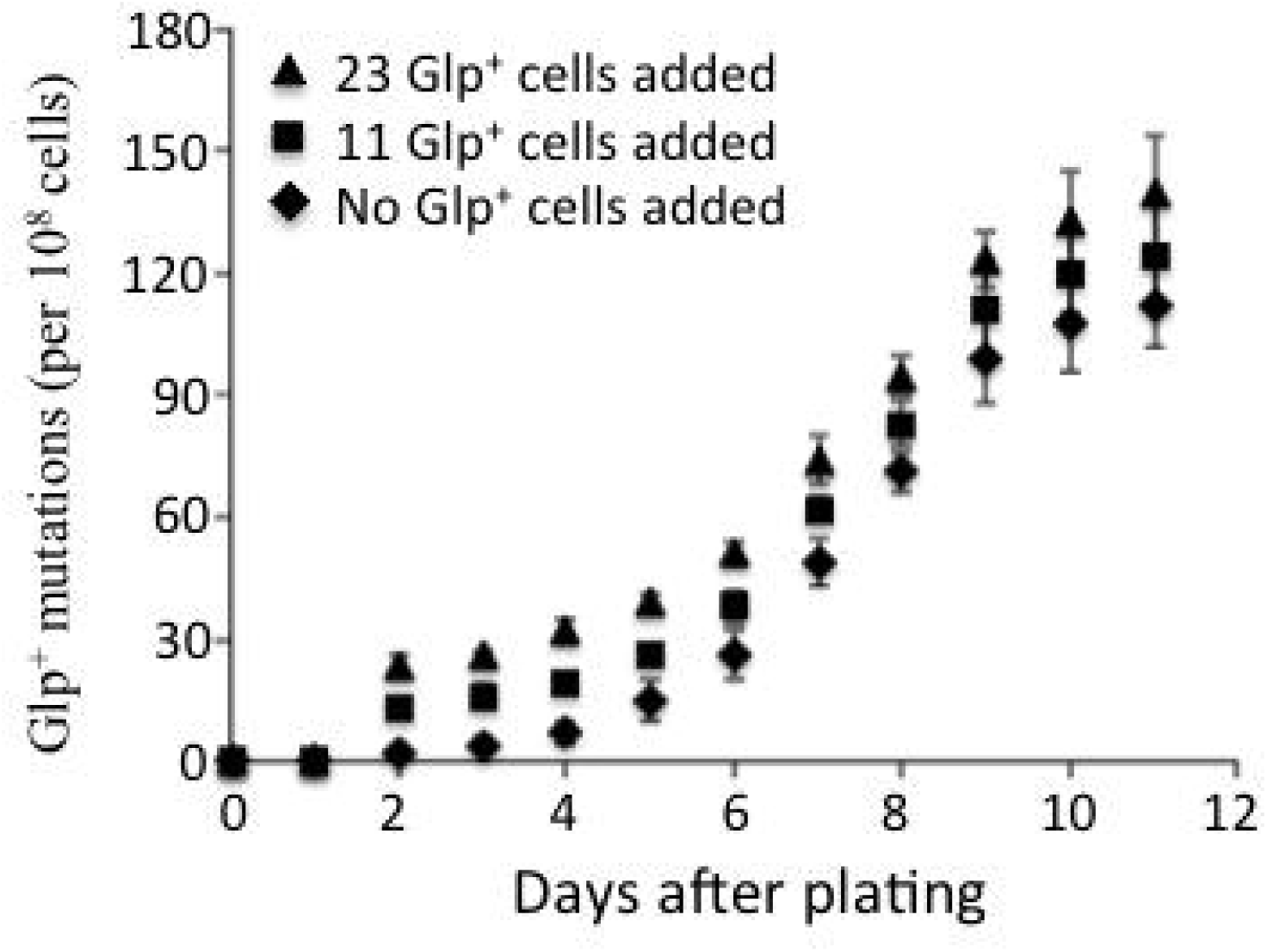
**Glp^+^ mutations in ΔcyaA cells on M9 + glycerol agar plates**. *AcyaA* cells (~10^8^) from a fresh LB culture were spread on M9 + glycerol (0.2%) agar plates. The plates were incubated at 30°C and examined for the appearance of Glp^+^ colonies (each colony represents an IS^5^ insertional mutation) at 24-h intervals. The mutation frequencies were determined as in Zhang and Saier (Zhang and Saier, 2009a). ♦ = no Glp^+^ cells added before initially plating; **■** and ▴ = 11 and 23 Glp^+^ cells were included before plating.

### IS^5^ activation of the glpFK operon in ΔcyaA cells is suppressed by exogenous cAMP

Exogenous cAMP can enter cells to increase the cytoplasmic concentration of this nucleotide (Saier et al., 1982). Figure 2A shows the effect of increasing concentrations of external cAMP on the *glpFK*-specific IS*5* insertional frequency in Δ*cyaA glpFK*_*cat* cells (see *Materials and Methods* and Supplementary Table 1). At a concentration of 10 µM, exogenous cAMP had only a slight inhibitory effect on IS*5* insertion, but at 100 µM, cAMP inhibited over 90%, whereas at 1 mM, cAMP essentially abolished IS*5* insertion (Figure 2A).

**Figure 2.**
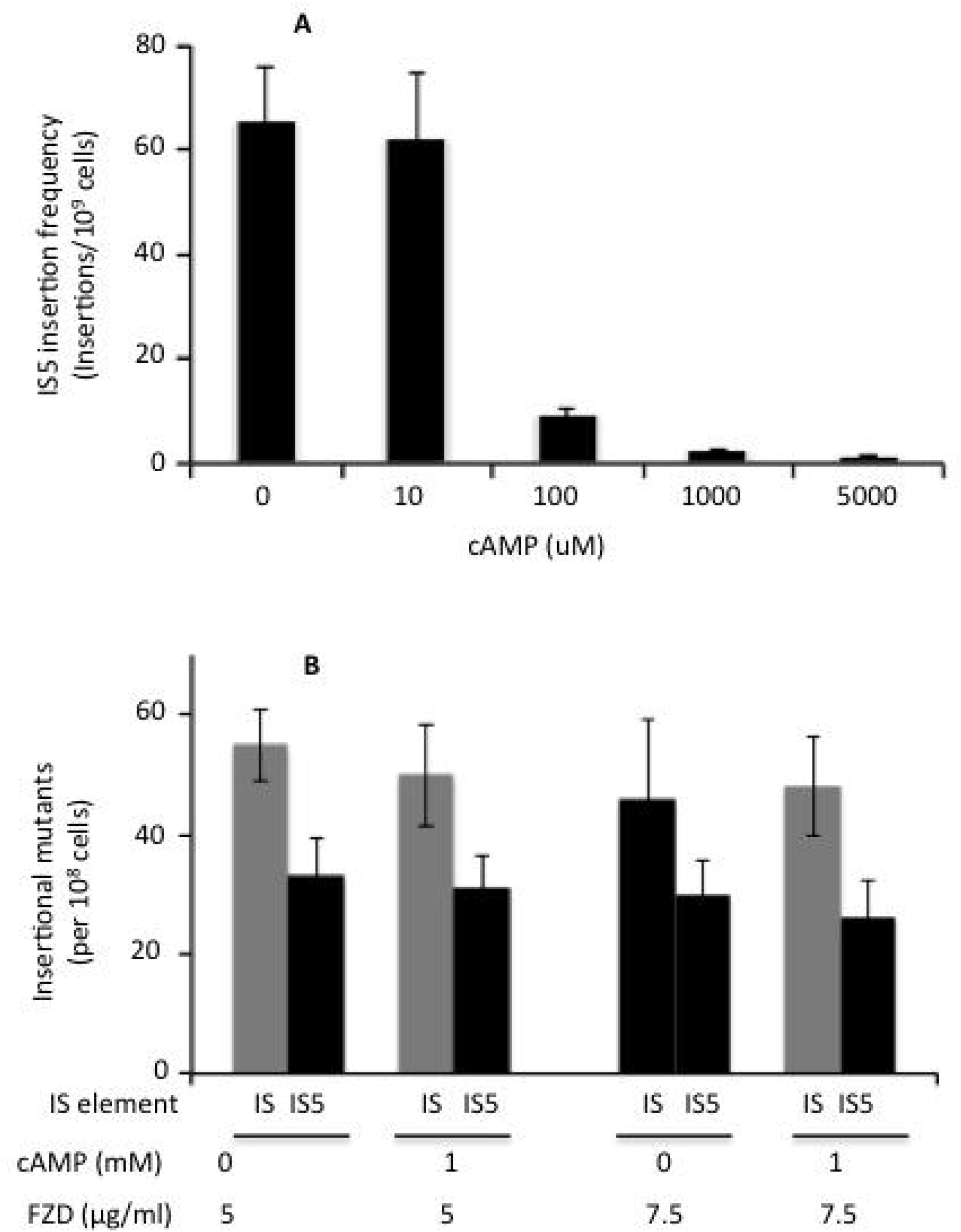
**Effects of cAMP on (A) IS^5^ insertion upstream of the *glpFK* regulatory region in *AcyaA* cells, and (B) in the *nfsB* gene as a control**. In A, a fresh LB culture from a single *ΔcyaA-Cat* (see Supplementary Table 1) colony was diluted 1000 x into 5 ml of LB ± cAMP (0 to 5 mM) in 30 ml glass tubes (2.5 cm x 20 cm). The tubes were shaken at 250 rpm in a 30 oC water bath shaker. After ~15 h, the cells were washed 1x (to remove residual cAMP) with carbon source-free M9 salts, serially diluted, and applied onto LB + glucose agar plates (for total population determination) and LB + glucose + Cm (16 ug/ml) agar plates (for IS^5^ insertional mutant population determination). In B, the cells (~2x 10) of a step I furazolidone (FZD) resistant (FZD^r^) mutant strain isolated from a *AcyaA-Cat* strain, were applied to nutrient broth (NB) agar plates, with furazolidone (5 or 7.5 jig/ml) ± cAMP (1mM). The plates were incubated at 30 oC for 36 h before being examined for the appearance of FZD^r^ mutants. Among these FZD^r^ mutants, IS^5^ or other IS insertional mutants in the *nfsB* gene were determined by PCR (see Materials and Methods). In all cases, the proportion of IS^5^ mutants (~60%) was the same.

To determine whether the decrease in IS*5* insertion frequency due to the presence of cAMP was specific to the *glpFK* promoter, we analyzed IS*5* insertion at the *nfsB* gene in Δ*cyaA* cells. Mutational inactivation of *nfsB* confers resistance to furazolidone (FZD) (Whiteway et al., 1998), and a significant fraction of inactivating mutations are due to IS*5* insertion (see *Materials and Methods*). As shown in Figure 2 B, cAMP did not influence the frequency of total insertional events (grey bars) or of IS^5^ insertional events (black bars) among FZD-resistant mutants within experimental error.

### The effect of cAMP on IS^5^ insertion frequency requires Crp-binding sites in the glpFK promoter

To determine whether the inhibition of IS*5* insertion upstream of the *glpFK* promoter by cAMP is due to the binding of the cAMP-Crp complex to the two adjacent Crp binding sites (*O_CrpI_* and *O_CrpII_*), present in the *glpFK* promoter, we analyzed the consequences of point mutations within these binding sites. These mutations essentially abolished the inhibitory effect of Crp on IS*5* insertion (Figure 3A). Although these Crp operator mutations eliminated binding of the cAMP-Crp complex, they did not change the *glpFK* promoter strength in the *cyaA* deletion background, and the promoter activity was still under the control of GlpR (Figure 3B). Thus, it can be concluded that inhibition by cAMP-Crp of IS*5* insertion into the *glpFK* activating site is due to the binding of the cAMP-Crp complex solely to these two operators present in the *glpFK* promoter region.

**Figure 3.**
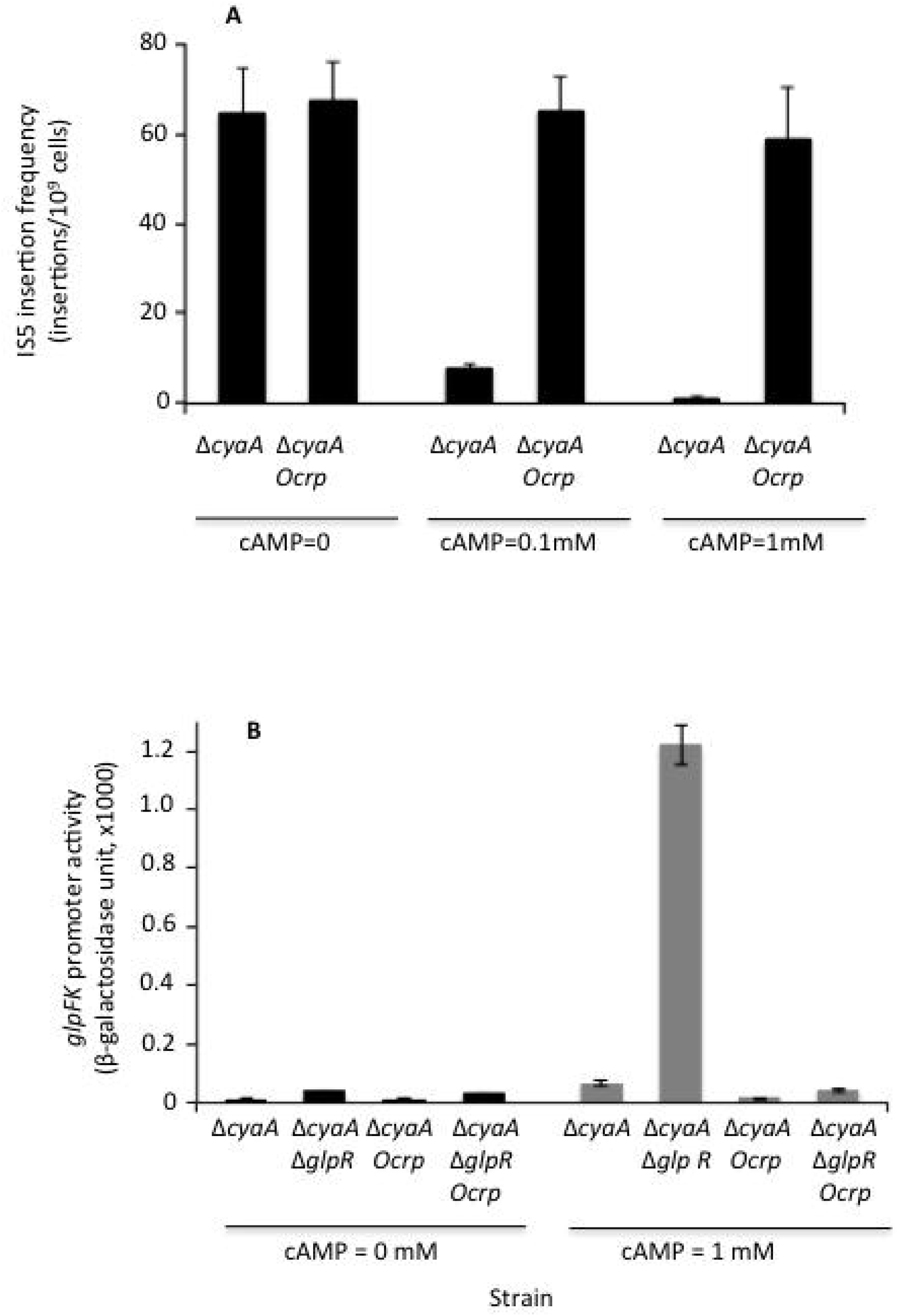
**Effect of Crp binding site mutations on IS^5^ insertion upstream of the *glpFK* promoter region (A) and the promoter activity in *AcyaA* cells grown in LB medium ± cAMP (B)**. The concentrations of cAMP are indicated below the x-axis. *O_crp_*= the mutated Crp binding sites (*O_crpI_* and *O_c rpII_*) in the upstream regulatory region of the *glpFK* operon to prevent Crp binding. In b, the activities of the *glpFK* promoter (P*glpFK*) and the same promoter (P*glpFK_O_crp_*) mutated in *O_crpI_* and *O_crpII_* were measured using the LacZ reporter in both Δ*cyaA* cells and Δ*cyaA* Δ*glpR* cells. See Supplementary Table 1 for the detailed strain information.

### Crp and GlpR independently affect IS^5^ insertion upstream of the glpFK promoter

To determine if GlpR plays a role in the inhibitory effect of Crp on IS*5* insertion, we deleted the *glpR* gene in the Δ*cyaA* background, yielding a Δ*cyaA* Δ*glpR* double mutant (Supplementary Table 1). Higher IS*5* insertional frequencies were observed in the Δ*cyaA* Δ*glpR* cells than in the Δ*cyaA* cells when grown in LB (no glycerol added) (Figure 4A). However, in the absence of GlpR, cAMP still exerted its inhibiting effect, presumably by binding to Crp, which then bound to its two *glpFK* operon binding sites, *O_CrpI_* and *O_CrpII_* (compare column 2 and column 4 in Figure 4A). Comparable inhibition was observed regardless of the presence of glycerol or GlpR. It was therefore concluded that regulation of IS*5* insertion by the cAMP-Crp complex occurs independently of glycerol and GlpR.

**Figure 4.**
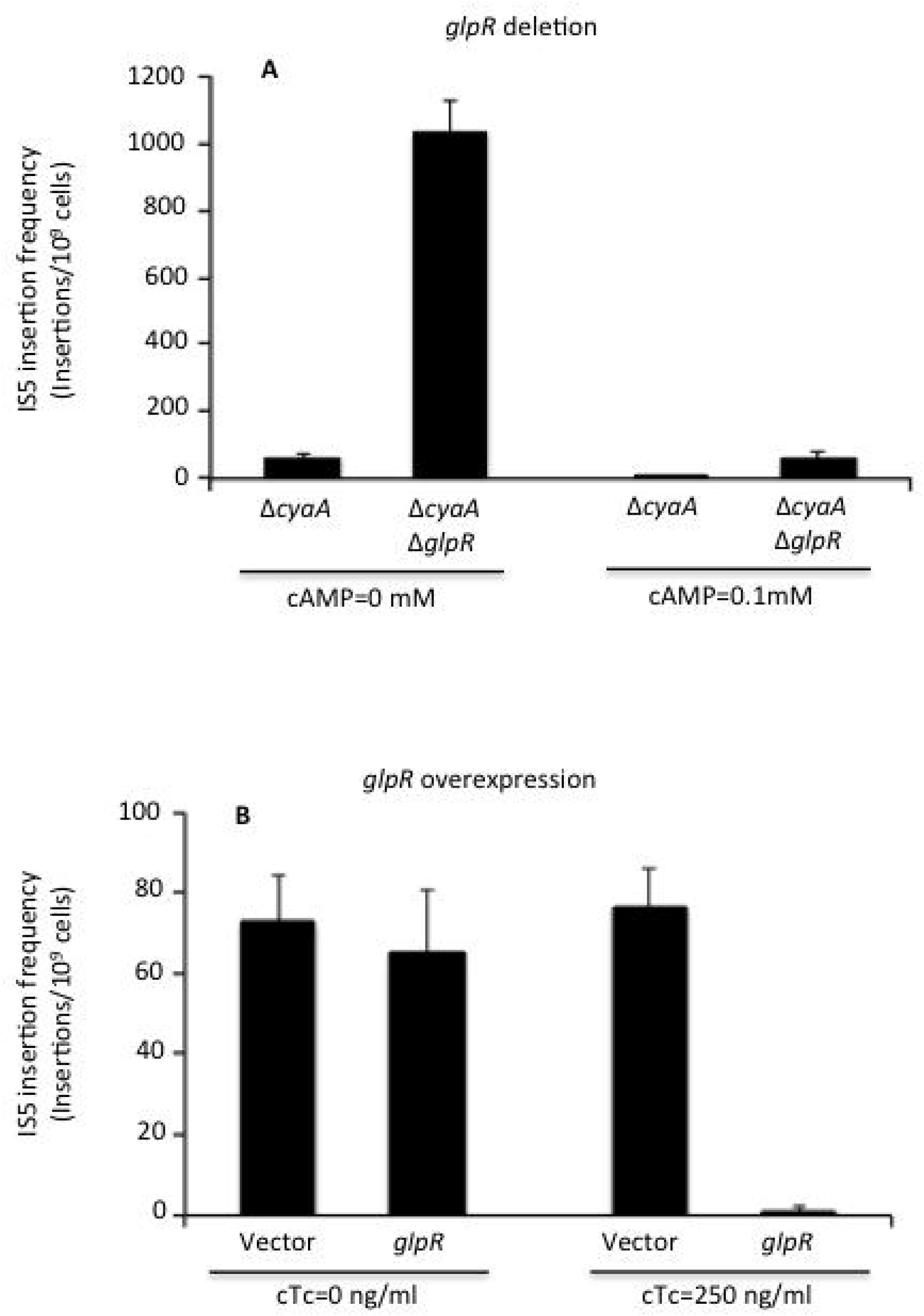
**Effects of *glpR* deletion (A) and *glpR* overexpression (B) on IS*5* insertion in the control region of the *glpFK* operon in** Δ***cyaA* cells grown in LB ± cAMP**. In A, the exogenous cAMP concentration was either 0 (left) or 0.1 mM (right). In B, vector: no *glpR* expression (control); GlpR indicates *glpR* expression; cTc = chloro-tetracycline (an inducer of the *tet* promoter) at the concentrations of 0 (left) or 250 ng/ml (right). *glpR* expression is under the control of the cTc induced promoter, P*tet*, in an expression vector. The Δ*cyaA* cells that constitutively produce TetR (repressing P*tet*) were used in these experiments. Note scale difference for Figure 4A and 4B.

When GlpR was over-produced in the absence of cAMP, GlpR still exerted its strong inhibitory effect (Figure 4B). In the left panel, the *glpR* gene was expressed at an extremely low level, and GlpR exerted only a minimal effect because chlorotetracycline (cTc), the inducer, was not present. When the cTc concentration was high (250 ng/ml), *glpR* was expressed at a high level, and the rate of IS*5* insertion into the *glpFK* upstream site was greatly reduced (see right panel of Figure 4b). The same experiments were conducted in the presence of cAMP (0.1mM). The IS*5* insertion frequency decreased, while overexpression of GlpR further inhibited IS*5* insertion (data not shown). These results suggest that GlpR and the cAMP-Crp complex exert their effects on IS*5* insertion independently of each other. These experiments also provide evidence that that GlpR and the cAMP-Crp complex can bind to the *glpFK* control region simultaneously.

### IS^5^ insertional activation of the glpFK operon occurs in a wild type background in the presence of 2-deoxyglucose

Since a reduction in cAMP promotes IS^5^ insertion in the *glpFK* activating site, we sought to determine whether environmental conditions that could lead to a reduction in cAMP concentrations could elevate IS*5* insertion into the *glpFK* promoter. Non-metabolizable glucose analogues, such as 2-deoxyglucose (2DG) and α-methylglucoside (αMG), (He and Liu, 2002, Holst and Williamson, 2004, Kumar et al., 2013, Moller, 2010, Tantanarat et al., 2012, Saier and Ballou, 1968, Xi et al., 2014) are known to lower cytoplasmic cAMP levels by inhibiting adenylate cyclase activity (Gabor et al., 2011, Gershanovich, 2003, Saier et al., 1996, Vastermark and Saier, 2014). These analogues also strongly inhibit growth on glycerol, at least in part due to inhibition of both cytoplasmic cAMP production by adenylate cyclase, and of cytoplasmic glycerol-3-phosphate (substrate/inducer) production by glycerol kinase (Kuroda et al., 2001, Peterkofsky et al., 2001, Schlegel et al., 2002, Saier and Reizer, 1994). We therefore asked if we could isolate IS*5* insertional mutants in a wild-type background on minimal M9 agar plates containing glycerol, inhibitory concentrations of 2DG or αMG and chloramphenicol. The results for αMG proved to be same as for 2DG, and consequently, only those obtained with 2DG are presented here.

In these experiments, a *glpFK-cat* (chloramphenicol acetyl transferase) fusion was used to measure IS*5* insertion (see Supplementary Table 1 and *Materials and Methods*). Regardless of the inhibitory glucose analogue used, 2DG/αMG, resistant (2DG^r^/ αMG^r^) mutants (which were also Glp^+^) could be isolated in a wild type *E. coli* genetic background (Jones-Mortimer and Kornberg, 1980, Kornberg et al., 2000, Rephaeli and Saier, 1980). As shown in Figure 5, PCR analyses of colonies that appeared after a five-day incubation at 30°C revealed two types of Glp^+^ mutants: (i) a majority that did not have an insertion in the *glpFK* promoter (dubbed “non-IS*5* mutants” here), and (ii) a minority (<10%; “IS^5^ mutants”) that had IS^5^ inserted into the glpFK promoter-activating site.

**Figure 5.**
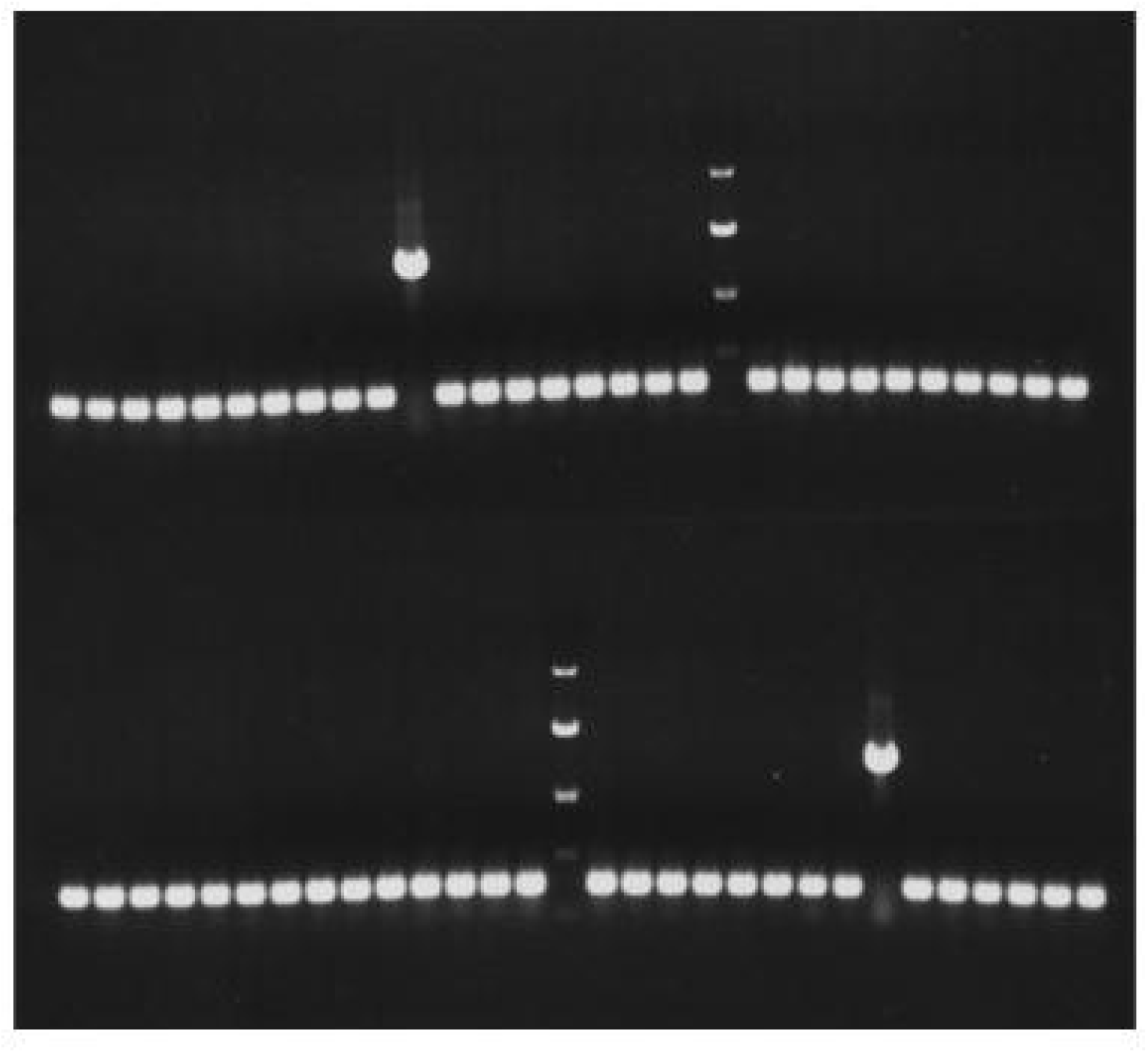
IS^5^ insertion upstream of the glpFK control region in wild type cells. Wild type cells carrying the glpFK-cat (chloroamphenicol acetyl transferase) fusion (See Methods section) were plated onto M9 + glycerol (0.2%) + 2DG (0.12%) + chloramphenicol (Cm) (55 μg/ml) agar plates. The plates were incubated at 30 oC. After 5 days of incubation, colonies were examined for the presence of IS^5^ upstream of the glpFK control region by PCR. In both the upper and the lower panels, a single IS^5^ insertion was obtained. A DNA marker showing four bands with known sizes (4 kb, 2 kb, 1 kb and 0.5 kb from above) is indicated in each panel.

Sequencing revealed that the non-IS^5^ insertion mutants did not have genetic alterations in the glpFK operon, or in the fruR gene [which appears to regulate crp gene expression (Zhang et al., 2014)]. Unlike IS^5^ insertional mutants, the non-IS^5^ mutants showed a pleiotropic phenotype in addition to their increased growth in a glycerol + 2DG medium, such as poor utilization of sorbitol (D-glucitol) and succinate (Table 1) which are utilized efficiently only when high cytoplasmic concentrations of the cAMP-Crp complex are available (Zhang et al., 2014).

**Table 1.** Growth of wild type, an IS^5^ insertional mutant and a non-IS^5^ mutant on minimal M9 agar plates supplemented with various carbon sources^1,2^.

| Strain | Glycerol | Sorbitol | Succinate | Xylose | Mannitol |
| --- | --- | --- | --- | --- | --- |
| Wild type | ++ | ++ | ++ | ++ | ++ |
| IS5 mutant | +++ | ++ | ++ | ++ | ++ |
| Non-IS5 mutant | ++ | + | + | ++ | ++ |

<sup>1</sup>Overnight LB cultures from single colonies were washed once with a carbon source-free M9 salt solution. The pellets were resuspended in the same M9 solution, and cells were inoculated onto the agar plates by streaking using a sterile transfer loop. The plates were incubated at 30 °C for 36 h before being examined for colony sizes.
<sup>2</sup>+, ++ and +++ denote the relative sizes of the colonies. All values are relative to growth of the wild type strain (++) on agar plates containing a single carbon source as indicated. Thus, +++ indicates increased size while + indicates decreased colony size.

To demonstrate that the Glp+ phenotype of IS^5^ mutants results solely from the IS^5^ insertional event, we carried out P1 transduction experiments to determine if the phenotypes of 2DGr, Cmr and Glp+, could be transferred together into another E. coli genetic background. In these experiments, we used two “wild type strains”, BW25113 and BW_Cat (Supplementary Table 1) (with or without the glpFK-cat fusion), as recipients. For both recipient strains, transductants were obtained using the IS^5^ insertional mutant (expressing the fused glpFK_cat operon) as donor, when plated on M9 + glycerol + 2DG ± Cm agar plates. The regulatory regions of 23 independently isolated transductants were amplified by PCR, and all were found to carry the IS^5^ element. DNA sequencing showed that IS^5^ was located in the same position and in the same orientation as described previously (Zhang and Saier, 2009a, Zhang and Saier, 2009c)). These results showed that in these mutants, IS^5^ insertion is necessary and sufficient to give rise to the 2DGr Glp+ phenotype.

### Short-term and long-term evolution experiments to evaluate if IS^5^-mediated activation of glpFK operon expression could have evolved in wild type cells

Initially, we conducted short term evolutionary experiments (several transfers) using the Δ*cyaA* strain in the presence of glycerol as sole carbon source (Figure 6A), glycerol plus 0.1 mM cAMP (Figure 6B), and glycerol plus 1.0 mM cAMP (Figure 6C). In the absence of cAMP, IS*5* insertional mutants appeared as the only species after 37 h of incubation with shaking. After several transfers, virtually 100% of the cells contained IS*5* in the *glpFK*-activating site (Figure 6A). When cAMP was added at 0.1 mM, IS*5* insertional mutants again appeared as the only species after a 61 h incubation with shaking. After the first transfer, virtually all cells contained IS*5* (Figure 6B). When exogenous cAMP was added at 1 mM, the appearance of these mutants was strongly inhibited. As a result, after the 6th transfer, only about 20% of the population were IS*5* insertional mutants (Figure 6C). These observations are consistent with the conclusion that cAMP-Crp complex inhibits IS*5* insertion upstream of the *glpFK* operon.

**Figure 6.**
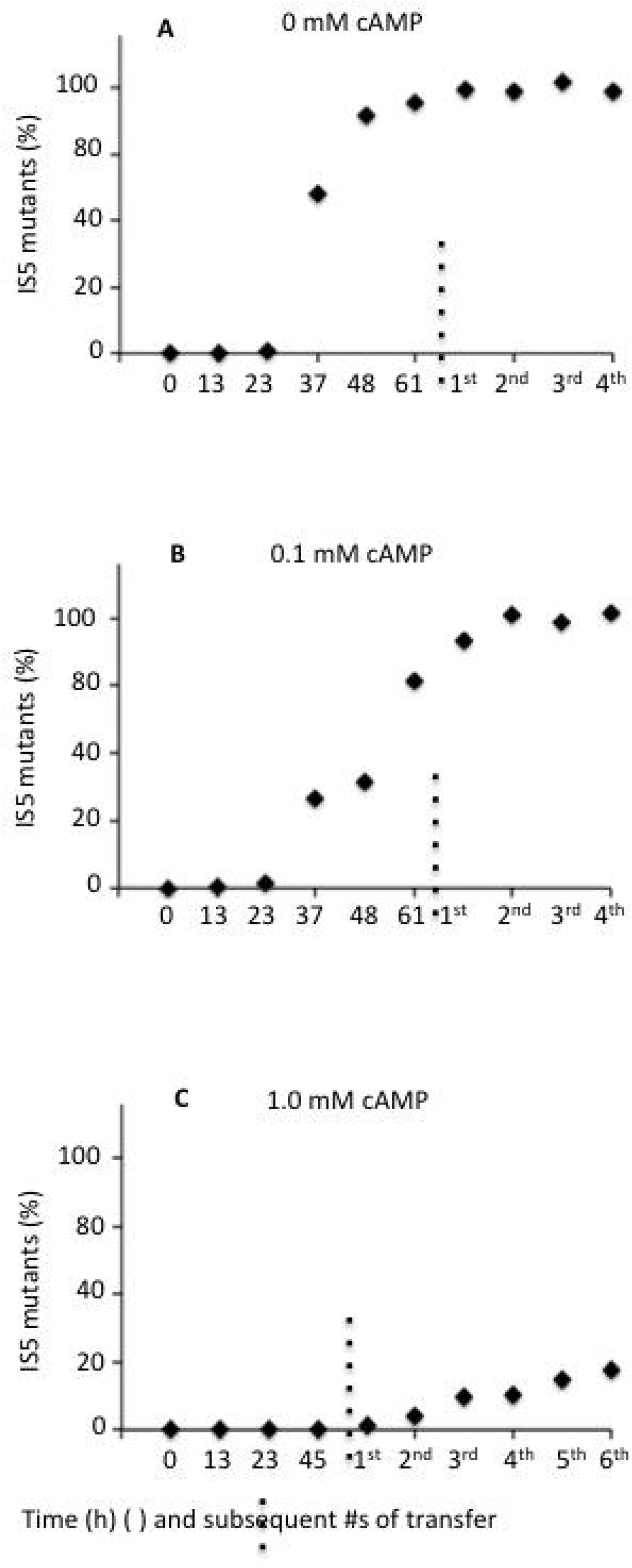
**Percentages of IS*5* mutant populations vs the total populations during growth in M9 + glycerol ± cAMP over time using strain** Δ***cyaA*_Cat**. For the first period [up to 61 h (A and B) or 45 h (C)], time is expressed in hours. Thereafter (vertical dotted line), time is expressed in numbers of transfers where each transfer took about a day. Figures A-C show the effects of increasing cAMP concentrations: A, 0; B, 0.1 mM; C, 1 mM.

In order to conduct long-term evolutionary experiments in a wild type background, we used *glpFK_cat* cells in which a chloramphenicol-resistance gene was transcriptionally fused to the *glpFK* operon in wild type cells such that *glpFK* activation by IS*5* insertion led simultaneously to a Glp^+^ as well as a Cm^r^ phenotype (see *Materials and Methods*). Prolonged incubation of *glpFK_cat* wild type cells under conditions analogous to those described above for the short-term Δ*cyaA* experiments revealed that, IS*5* mutants became an appreciable fraction of the population after several generations in minimal glycerol medium with 2-deoxyglucose, with (Figure 7A) or without (Figure 7B) chloramphenicol. When chloramphenicol was present, IS*5* insertional mutants first appeared after six transfers, and then continued to accumulate during the remainder of the experiment (17 transfers) at the end of which about 25% of the cells bore the IS^5^ insertion (Figure 7A). When chloramphenicol was absent, it took a little longer; insertional mutants became appreciable following the eighth transfer, and accounted for over 20% of the cells after 20 transfers (Figure 7B). These experiments show that *glpFK* activation by IS*5* is advantageous when wild type cells are exposed to glycerol in the presence of a sugar analogue inhibitor of adenylate cyclase such as 2-deoxyglucose (Novotny et al., 1985). Thus, the process we had previously described in Δ*crp* and Δ*cyaA* mutants could have evolved in wild type cells in response to specific environmental conditions.

**Figure 7.**
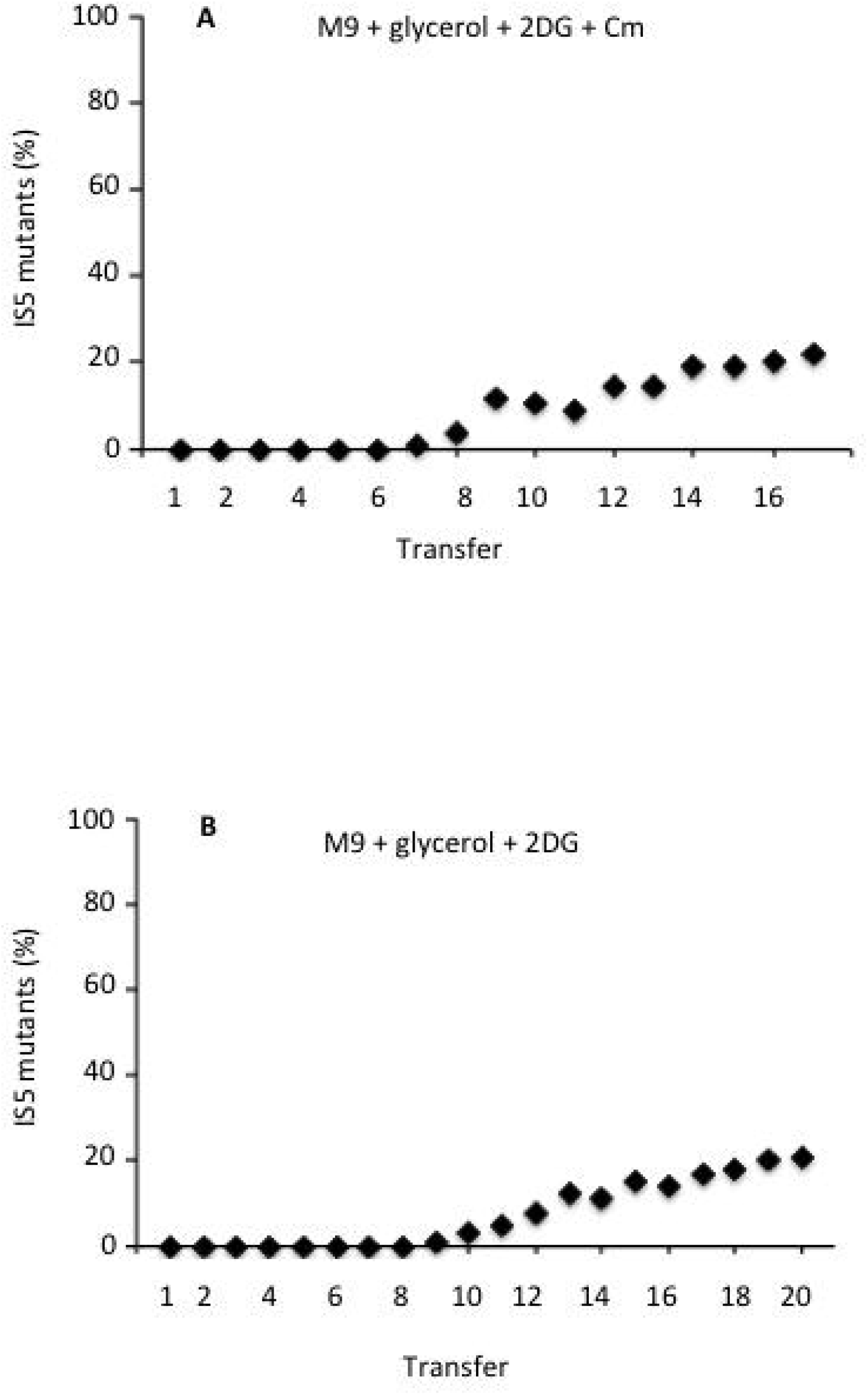
**Percentages of IS^5^ insertional mutant populations versus total mutant populations over time using the “wild type” strain carrying the chromosomal *glpFK*-*cat* fusion**. Cells were incubated with shaking at 30 ^o^C in 5 ml of M9 + glycerol (0.2%) + 2DG (0.13%) plus (A) or minus (B) chloramphenicol (Cm) (60 μg/ml) in glass tubes (20 mm x 200 mm). After 2.5 days of incubation, 5 μl cultures were transferred to 5ml of the same media (1^st^ transfer) and incubated with shaking at 30 ^o^C. Later, every two days, 5 μl cultures were transferred to new media and grown under the same conditions. For every transfer, the cultures were diluted with carbon source-free M9 salt solution and plated onto M9 + glycerol + 2DG + Cm agar plates. After 2 days of incubation at 30 ^o^C, 100 colonies were examined by PCR for the presence of IS*5* in the *glpFK* control region (See Figure 5).

## Discussion

Transposons (transposable elements; jumping genes) were first described by Barbara McClintock (McClintock, 1950), recipient of the 1983 Nobel Prize (Kenez, 1984, van de Putte, 1983). Initially identified in corn, transposable elements occur in virtually all living organisms, where they cause mutations that can inactivate genes, activate genes, and mediate chromosomal rearrangements (Zhang and Saier, 2011). Different types of transposons (especially retrotransposons) comprise about 30% of the human genome (Cordaux and Batzer, 2009). Why are they ubiquitous, and what are their primary functions? What is responsible for their broad occurrence in genomes of all types of living organisms? The answers to these questions are not yet in hand, but it is clear that transposons contribute to genomic instability and genetic innovation (Huang et al., 2012).

Long-terminal-repeat (LTR) retrotransposons are known to jump around within eukaryotic genomes (Curcio et al., 2015). To avoid damaging resident genes, they have been selected to integrate away from protein-coding sequences. For instance, the fission yeast LTR retrotransposon, Tf1, inserts at nucleosome-free regions in gene promoters. Jacobs et al. (Jacobs et al., 2015) recently showed that Tf1 is directed to these insertion sites by specific DNA binding proteins such as Sap1, a situation that parallels in some respects the observations reported here. Thus, regulation of transposon insertion by DNA-binding proteins may prove to be common in Nature, perhaps even ubiquitous.

Adaptive mutation, the genome-wide acceleration of mutation rate under stress conditions, is well-documented, and in some cases, the mechanisms involved are understood (Shee et al., 2011). An understanding of the basis for mutations that increase fitness, give rise to drug resistance, and enable the emergence of pathological traits in microorganisms, are also relevant to human, animal and plant health, at both the individual and the population levels (Gordo et al., 2011).

Over the last several decades, a number of mechanisms that accelerate mutagenesis under stressful conditions have been described. Among these have been those that activate cyptic operons by transposon insertion under starvation conditions where such activation is beneficial to the cell (Hall, 1999). Another example is provided by enhanced IS insertional mutations causing metal resistance in bacterial cells subjected to toxic levels of zinc (Vandecraen et al., 2016). The activation of the *glpFK* operon by IS*5* insertion represents an adaptive mutagenic mechanism where some, but not all, molecular mechanisms leading to the activation of the operon are understood (Zhang and Saier, 2009b, Zhang and Saier, 2009d). In this paper we advance the state of knowledge of this system by showing that transposition into the upstream promoter region of *glpFK* is controlled independently by two defined DNA binding proteins (GlpR and the Crp-cAMP-Crp complex). We show that IS*5* insertion into the *glpFK* promoter can be regulated by exposure to environmental toxic sugar analogues, and propose that such exposure may constitute a selective environment that favored the evolution of a gene activation mechanism that requires a specific transposition event.

In our experiments with wild type *E. coli* cells, we used 2-deoxyglucose (2DG) as described in this report or methyl α-glucoside (αMG) (data not presented) to lower cytoplasmic cAMP levels and to inhibit glycerol utilization (Saier, 1998, Saier, 1989, Saier, 2001, Saier et al., 1995). But are comparable non-metabolizable glucose analogues found in Nature? In fact, numerous toxic and non-metabolizable sugar analogues are synthesized by microorganisms, plants, fungi and man. They include deoxy sugars such as 2-deoxyglucose, methylated sugars, such as 3- and 6-0 methyl glucose, fluoro sugars and a variety of α- and β-glycosides such as methyl α-glucoside (He and Liu, 2002, Holst and Williamson, 2004, Kumar et al., 2013, Moller, 2010, Tantanarat et al., 2012, Saier and Ballou, 1968, Xi et al., 2014). Thus, the conditions that promote IS*5* hopping into the *glpFK*-activating site could have evolved under the pressures of natural selection. Supporting a possible evolutionary role for transposition under starvation conditions, it was recently reported that prolonged starvation leads to significantly elevated expression of transposases in *E. coli*, including the *insA* gene, encoding the IS*5* transposase (Arunasri et al., 2014).

## Concluding Remarks

We have shown that the IS*5*-mediated *glpFK*-activating insertional site is a conditional transposition hotspot. It is an insertional hotspot in low cyclic AMP and high glycerol, but insertion at this site is essentially undetectable in the presence of other carbon sources, or high levels of cyclic AMP in the absence of glycerol. The experimentation described in this report explains the effect of cAMP for the first time, and provides plausible conditions under which such a mutational mechanism could have evolved in a wild type genetic background. Thus, our results demonstrate that the activated cAMP-Crp complex strongly inhibits IS^5^ hoping into the *glpFK* activating site when bound to its DNA-binding sites in the upstream *glpFK*-promoter region. Our results also showed definitively that in the lab, one can isolate these activating mutants in a wild type genetic background in the presence of glycerol and inhibitory concentrations of 2DG or α-MG that represent adverse environmental conditions.

## Acknowledgements

We thank Fengyi Tang, Joshua Asiaban, Sabrina Phan and Yongxin Hu for assistance with manuscript preparation, and Chika Kukita and Robert Palido for technical assistance with some of the experiments. This work was supported by NIH grants GM109895 and GM077402.

